# Programmatic modeling for biological systems

**DOI:** 10.1101/2021.02.26.433125

**Authors:** Alexander L.R. Lubbock, Carlos F. Lopez

**Affiliations:** Department of Biochemistry, Vanderbilt University, Nashville, Tennessee 37212, United States of America; Vanderbilt-Ingram Cancer Center, Vanderbilt University, Nashville Tennessee 37212, United States of America; Department of Biomedical Informatics, Vanderbilt University, Nashville, Tennessee 37212, United States of America

## Abstract

Computational modeling has become an established technique to encode mathematical representations of cellular processes and gain mechanistic insights that drive testable predictions. These models are often constructed using graphical user interfaces or domain-specific languages, with SBML used for interchange. Models are typically simulated, calibrated, and analyzed either within a single application, or using import and export from various tools. Here, we describe a programmatic modeling paradigm, in which modeling is augmented with best practices from software engineering. We focus on Python - a popular, user-friendly programming language with a large scientific package ecosystem. Models themselves can be encoded as programs, adding benefits such as modularity, testing, and automated documentation generators while still being exportable to SBML. Automated version control and testing ensures models and their modules have expected properties and behavior. Programmatic modeling is a key technology to enable collaborative model development and enhance dissemination, transparency, and reproducibility.

**Highlights:**

- Programmatic modeling combines computational modeling with software engineering best practices.
- An executable model enables users to leverage all available resources from the language.
- Community benefits include improved collaboration, reusability, and reproducibility.
- Python has multiple modeling frameworks with a broad, active scientific ecosystem.

## Introduction

Mathematical modeling of cellular processes for mechanism exploration has now become commonplace using various techniques [1–5], but challenges remain as to how models should be built, calibrated, analyzed and interpreted to extract much-needed mechanistic knowledge from experimental data. Historically, methods and techniques from other fields have been directly imported to systems biology with varying success. For example, early interpretations of cellular processes as circuits provided insights about basic regulatory motifs that could explain cellular behaviors [6]. Similarly, techniques from chemistry, physics, and various engineering disciplines have been used to model cellular processes [7,8], but due to the spatiotemporal complexity of cellular processes, from femtosecond/nanometer electron transfer reactions to years and meter scales in tumor growth, no established paradigm has emerged to capture the full complexity of cellular processes. Multiple tools have been developed to achieve specific modeling tasks. For example, COPASI [5], RuleMonkey [9], Simmune [10], and StochSS [11] all provide graphical user interfaces that cater to non-expert modelers wishing to encode mechanistic representations of biological processes. More abstract approaches such as BioNetGen [12], Kappa [13], and CobraPy [14] employ a domain-specific language (DSL) to describe and simulate models. However, most tools are self-contained platforms with a small set of included methods and analyses, limiting access to other standalone simulation tools such as StochKit [4], SciML tools[15], URDME [16], SmolDyn [17]. Similarly, optimization techniques ranging from vector-based optimization methods [18,19] to probabilistic-based methods [20*,21] exist in yet another isolated domain. Therefore, the current modeling and simulation ecosystem is compartmentalized and fractured, and thus, unification and intercompatibility efforts are sorely needed.

Valuable efforts toward unification have been put forth to create standards for model instantiation, simulation, analysis and dissemination [22,23**,24–26]. Of these, Systems Biology Markup Language (SBML) is perhaps the most successful to date. However, mathematical modeling for cell biology remains challenging to scale - both vertically (larger, more complex models) and horizontally (more active collaborators). While mathematical tools are the obvious way forward to describe cellular processes, the complexity challenge results in a knowledge base that is highly domain specific, with some notable exceptions [27*].

A novel, more flexible approach to encode knowledge about biological processes as computer programs is slowly emerging and gaining momentum [3,28,29]. In this approach, biological models are no longer static documents, but computer code that aggregates community knowledge and opens doors toward crowd-driven mathematical models of biological processes. Although computer languages like Lisp [30] and proprietary packages such as MATLAB have been used toward this goal, we believe Python provides the largest ecosystem, myriad learning resources, and large applicable base to unify modeling practices in the field. Adopting a programmatic modeling paradigm for systems biology automatically accrues decades of computer science practices including structured documentation, integrated development environments (IDEs), (model) version control, code-sharing platforms, code testing frameworks, and importantly, literate programming/computational notebook dissemination. Here, we review the recent developments in programming-based approaches for systems biology. The structure of the manuscript is motivated by the model specification, simulation, calibration, analysis, and visualization paradigm/pipeline, commonly practiced in systems biology. Throughout, we note how this approach could be supplemented and improved by incorporating best practices from software engineering (Figure 1).

**Figure 1:**
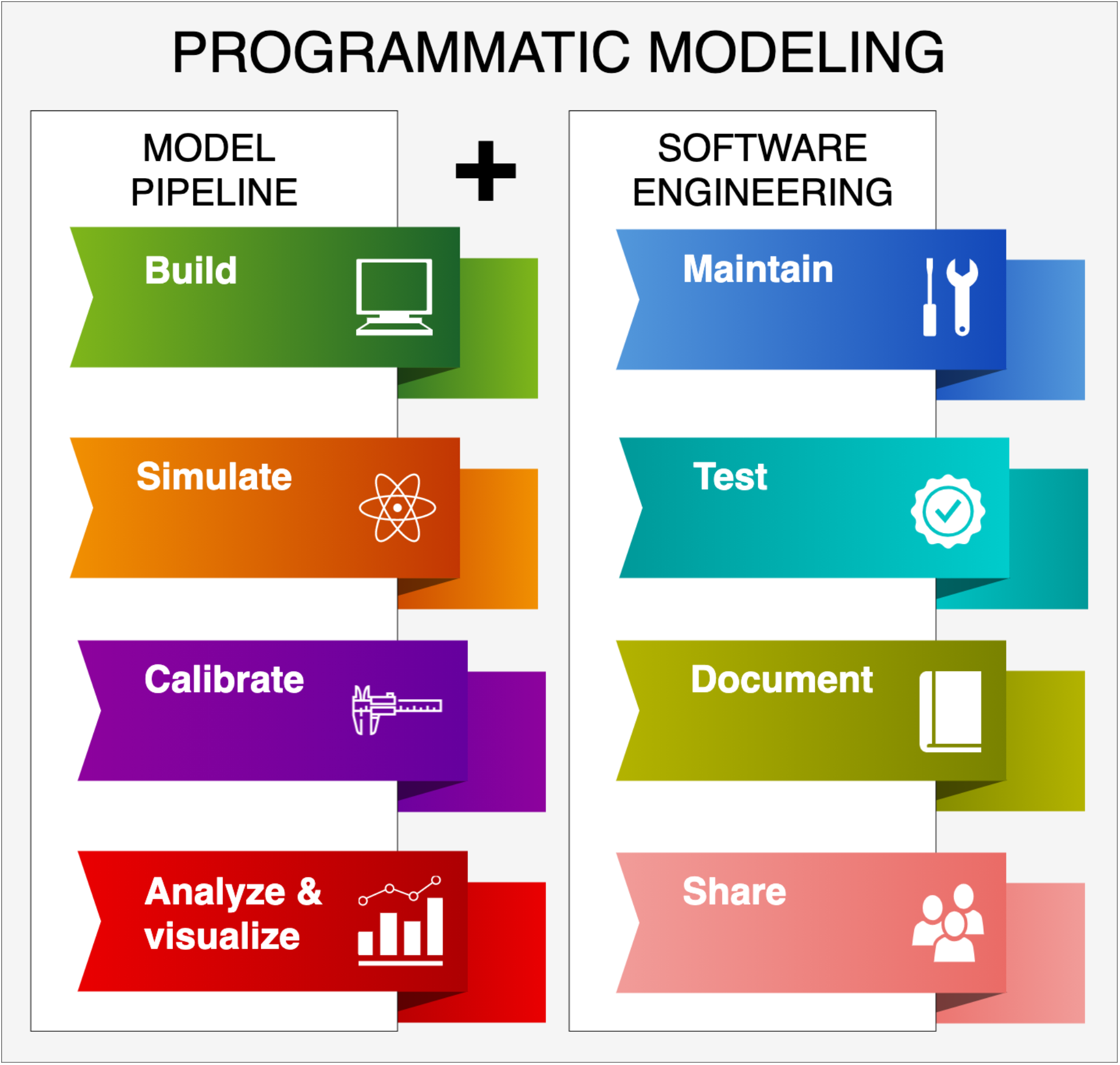
The traditional modeling paradigm in systems biology entails model building, simulation, calibration, and analysis (left column), which is carried out with myriad tools and practices. Software engineering practices can add much needed structure to the practice through maintenance, testing, documentation and sharing paradigms (right column), vetted by a the software community.

## Model specification

Traditionally, encoding a model of biochemical reactions would require the user to write each equation by hand, encode these into a solver, and run the simulations [31]. Although this is still common practice for smaller model systems, these lists of equations often lead to a model dead-end as the biological context is completely lost in the mathematical representation, which hinders model reuse. Reaction-based modeling formats add one layer of abstraction where the user instead writes chemical reactions of the form *A* + *B* ↔ *C* in a program-specific notation and the computer parses this information into a mathematical representation [32]. These DSLs can operate either through a GUI that generates the code in the background, or directly through a text editor. For example, Antimony [32] requires manual enumeration of every species and reaction explicitly. However, signaling pathways often comprise a large number of molecular complexes, which can assemble in multiple orders, leading to a large number of reactions and intermediate species during complex assembly and degradation. Therefore, traditional approaches become unwieldy as model systems become larger, learning to model dead-ends. Another level of abstraction is presented by rule-based modeling formalisms whereby *reaction rules* rather than explicit reactions (or equations) are used to encode the system [3,12,13]. A reaction rule is a template for reaction patterns to be enumerated and instantiated recursively, starting from a defined list of initial species, thereby saving the user time and reducing error-prone repetition. In rule-based approaches, the reaction center (the relevant molecular components for a given reaction) is separated from the context (attached molecular components which have minimal or no effect on the reaction). These approaches often require a pre-processing step to generate the network of nodes (chemical species) and edges (chemical reactions) from the initial pool of chemical species and a set of reaction rules. However, network-free methodologies have been proposed to bypass the network generation step [33].

Model specification can also be embedded into General Purpose Programming Languages (GPPL) to provide a more powerful approach to biological modeling. In the programmatic modeling paradigm, the model is encoded as an executable piece of code, thereby offering all the advantages of a full-fledged computer programming language (Figure 2). Modularity, in which a model can be split into smaller, reusable code objects, is perhaps the most useful aspect for cell biology modeling. For example, PySB currently includes a library of 25 macros (small modules or functions) that encode reaction patterns commonly found in biology such as catalytic activation, molecule-molecule inhibition, or complex oligomerization. From a user perspective, GPPLs have greater integration with IDEs than DSLs, thus allowing syntax highlighting and checking, and navigation between functions. The model is also inspectable at runtime, allowing searching and filtering of model components. For example, a user could check whether certain species or reactions are present before simulation commences. Currently, the most used modeling frameworks using the programmatic modeling paradigm in Python are PySB [3], written in and using Python, and Tellurium [29], which is written in Python but uses Antimony [32], a DSL with function support, for model specification.

**Figure 2:**
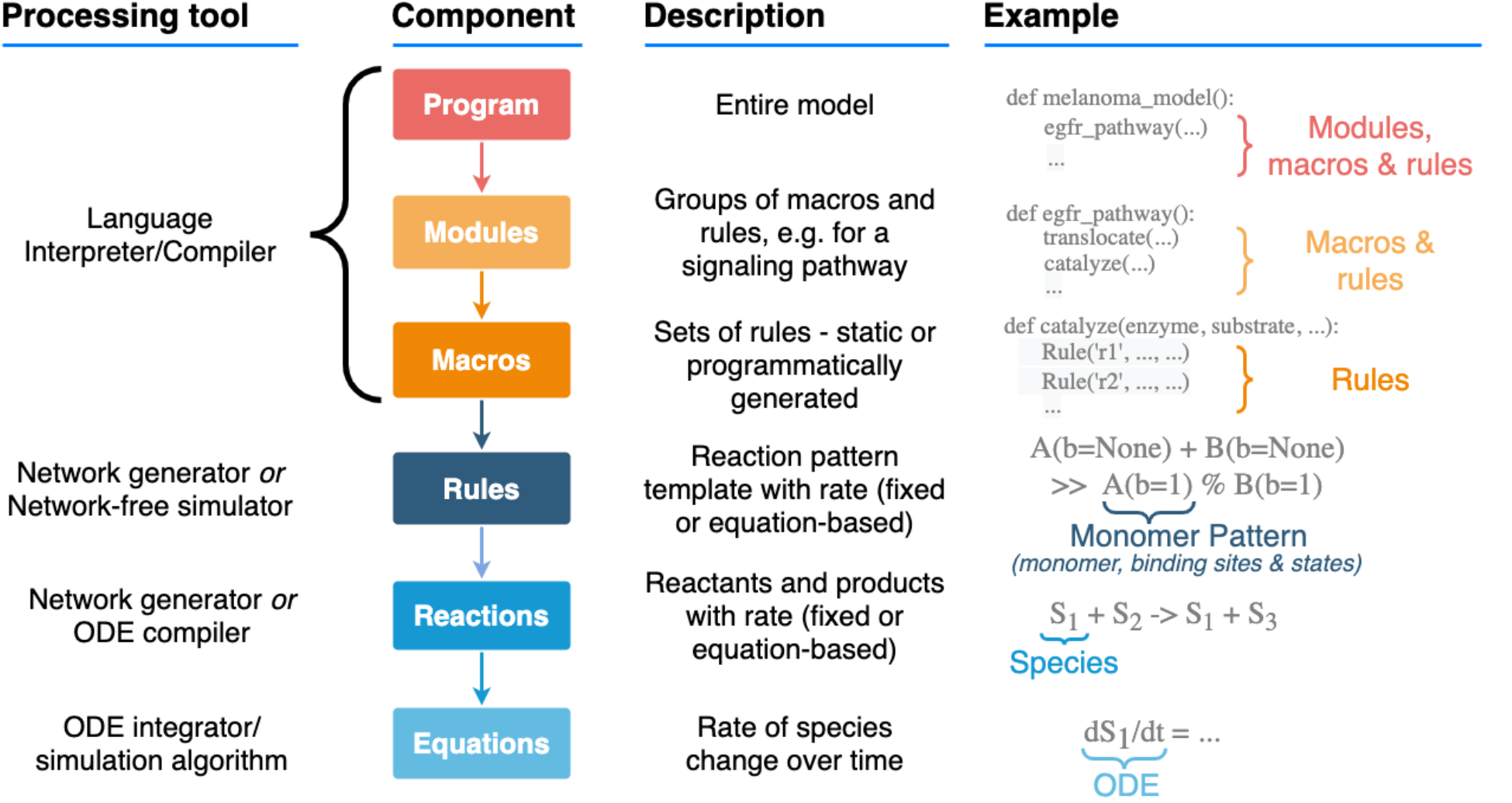
Levels of abstraction in programmatic modeling. Models are composed of modules and macros, which are handled by the programming language interpreter/compiler; rules encode sets of reactions using structured pattern templates; reactions specify biochemical species’ transformations; and finally equations are handled by an ODE integrator or simulation algorithm directly.

## Model simulation

Model simulation involves numerically solving the model equations to obtain trajectories for dynamically controlled species. Concentrations or molecule counts of chemical species in the model are the most commonly simulated quantities. Integration of systems of ordinary differential equations (ODEs) for deterministic simulations is the most common model simulation approach. Many ODE integrators are available and the best choice depends on model stiffness, desired integrator tolerances, and other requirements. In Python, a family of integrators is available through SciPy [34**] including VODE and LSODA, but many other solvers have been proposed. Other commonly used solver suites include StochKit (Stochastic Simulation Algorithm) [4,35], BioNetGen (CVODE, SSA, tau-leaping algorithm, partition-leaping algorithm) [12], cupSODA (GPU ODE) [36], GPU_SSA (GPU SSA) [37], and Libroadrunner (CVODE, SSA) [38]. Within the Python ecosystem, PySB provides a simulation class that enables users to use many of these simulation tools or to connect new tools as needed. In addition, users of other Python-based tools such as Tellurium can also take advantage of these resources.

## Model calibration

Model calibration is the process of adjusting model parameters to match experimental data, also known as parameter estimation/optimization when applied to parametric models. The most common form of model calibration involves a process of running many simulations (thousands to millions or more) and checking the distance between model and experimental data error using an objective function, which gives a measure of the model’s simulation “error” versus experiment; for a review see [39]. Since dynamic data for signaling models are hard to come by, the modeler often only has data for a few species, and thus model calibration often leaves a model underdetermined - multiple parameter sets fit the data equally well [40]. The concept of parameter “sloppiness” states that only a few “stiff” combinations of parameters are important in determining model outcomes, and others are “sloppy” and have little effect. Thus, an undetermined model can still be useful in predicting biological properties [41]. However, the interpretation of large, underdetermined models in the context of limited data is still up for debate. Lessons from e.g. hydrology and climate modeling have been highly influential toward addressing these issues [20*,42,43].

The landscape of model parameters is often envisioned as a multidimensional surface with “height” representing the objective function, where the (ideally global) minimum or minima (representing the best fit(s)) must be found. SciPy [34**], for example, includes gradient descent and simplex-based methods. However, the curse of dimensionality means that local optimization can give far-from-globally optimal results as the number of model parameters increases. Finding the global minimum of a multivariate nonlinear model is NP-hard [44], however several methods can make statistically good approximations. Markov Chain Monte Carlo sampling methods are among the most popular algorithms [45]. General purpose optimization toolkits for Python include SciPy.optimize [34**] and Pyomo [46]. We have found that DEAP [19] provides excellent support for PSO and genetic algorithm-based optimization.

Given the dearth of data available for biological model calibration, conditional probability (Bayesian) approaches are gaining traction. These approaches provide a probabilistic interpretation of model parameters [47], including uncertainty quantification, at the cost of increased computer time. However, new GPU-based integrators mitigate this problem. Excellent tools for Bayesian parameter inference include PyDREAM (which can readily take PySB models) [20*], PyBioNetFit [48*], PyPESTO [49], PyMC3 [50], and PySTAN [51], although popular data-science tools such as TensorFlow [52] and PyTorch [53] also provide Bayesian inference capabilities. ABC-SysBio [54] provides a hybrid solution to the computation problem but still within a Bayesian context.

## Model analysis and visualization

Model analysis and visualization is likely the least developed area in systems biology as no clear standards have been proposed. In general, modelers explore the chemical species concentration trajectories in their model to infer mechanistic behaviors and properties. Exploration of biochemical flux through reactions is highly challenging with some notable attempts toward this goal in the literature [7,47], but much work is still needed. For visualization, perhaps the most useful tool in Python is matplotlib [55], which provides flexible graphing capabilities. Other Python tools include Seaborn (https://seaborn.pydata.org/), Plotly [56], and Mayavi [57]. Network visualization is perhaps the other major area of model analysis that is addressed in various ways in Python. For example, PyVIPR [58*] is a visualization tool built on Cytoscape.js [59] for rule- and reaction-based models which animates model dynamics over time, overlaid on a graph. MASSPy [60] also provides some visualization capabilities for metabolic models. We note, however, that excellent tools for graph manipulation in Python exist, such as NetworkX [61].

## Model sharing and modification

Perhaps the most appealing benefit for the systems biology community from program-based paradigm is the use of literate programming for model and results dissemination. Introduced by Donald Knuth, literate programming is a paradigm whereby the code and the document coexist in an interactive format [62]. Jupyter Notebook, a popular format, has been described as “data scientists’ computational notebook of choice” [63]. Jupyter Notebooks allow analyses to be run in a web browser, checked into version control, and include documentation alongside analyses, in turn improving transparency and reproducibility. We believe that Jupyter notebooks are a highly desirable step forward in systems biology as it greatly contributes to model transparency, revision, and dissemination, and should be included in paper submissions where computational simulation and analysis are involved.

Programmatic models’ code can be managed using existing version control tools. Git has emerged as the *de facto* standard for version control, providing powerful capabilities for decentralized editing, branching, and merging, with online platforms such as GitHub adding a collaborative interface for change management, commenting, and other functions. In PySB, models are Python programs, and so can be imported like other Python modules and extended or modified. The code can be inspected, for example the model can be searched for species or reactions using pattern matching. Tellurium’s antimony language has an import function, but previous model definitions are currently not programmatically searchable or modifiable.

Good documentation can be vital to ensure model reproducibility and interpretability by others. Sphinx (sphinx-doc.org) is a *de facto* documentation standard for Python code, which allows code comments to be compiled into multiple formats including website (HTML) and PDF. The former can be combined with continuous integration, for always up-to-date documentation (readthedocs.io).

## Model checking and testing

Complex biochemical models present challenges in both ensuring they are correctly encoded, and ensuring their dynamics meet a given specification. In software engineering, it has become common practice to build an accompanying test suite while developing code, which runs the code under scrutiny to test that works as expected. Subtle errors can be introduced as models grow larger. In our opinion, the field should establish minimum standards to ensure software is runnable, reproducible, and meets basic quality standards [64]. In the context of models-as-programs, unit and integration tests can be borrowed from software engineering to ensure code correctness. Unit tests refers to code which checks the functionality of other, minimal units of code; integration tests check that units work as expected when combined. Python has several frameworks for testing, PyTests is a popular option with a plugin for Jupyter Notebooks [65]. PySB introduces a framework for testing static properties of rule-based models after network generation; for example, checking that certain species are produced by the reaction network, or that certain reactions are present. Using continuous integration (CI), these tests can be run automatically when changes are made and checked into version control, and/or on a regular basis. Running tests regularly is recommended because, even if a model itself does not change, changes to software dependencies could lead to unexpected errors. The importance of this is emphasized by a recent review, which found a majority of Jupyter Notebooks were not automatically reproducible, often due to dependency errors [66**]. For open-source models, these tests can be run for free using services such as Github Actions, Travis, and Circle CI. Finally, we recommend containerization technologies such as Docker [67] and Singularity [68], which bundle model and software dependencies together in a self-contained environment, further aiding reproducibility and cross-platform compatibility.

## Conclusions

Python has recently turned 30 years old and is now one of the most popular programming languages in the world. There are many reasons for its success, but a key insight of its creator is that code is read much more often than it’s written [69]. The same principle applies to models, which emphasizes the importance of clear documentation, transparency of approach, and the separation of model specification from simulation and downstream analysis code. These efforts are central to improving reproducibility, code maintenance, and model extensions, by original authors and third parties.

For beginners interested in modeling cell signaling, we recommend either the PySB or Tellurium frameworks, both of which have high quality documentation and active communities for support. We expect the Python modeling ecosystem will continue to grow, and efforts for framework and package interoperability to increase.

**Table 1:** List of key frameworks, tools, and services for programmatic modeling in Python. BSD, MIT, and PSF are permissive software licenses. GPL is a “copyleft” software license. *Free for open-source projects.

| Tool or service | Usage terms | Notes |
| --- | --- | --- |
| **Frameworks** |  |  |
| PySB [3] | BSD | Bespoke Python object-based model format, multiple simulation backends |
| Tellurium [29] | Apache | Bespoke DSL (Antimony), ODE and SSA simulation backends |
| PySCeS [70] | BSD | Bespoke DSL, ODE simulation backend |
| ScrumPy [71] | GPL | Metabolic modeling |
| CobraPy [14] | GPL | Metabolic modeling |
| **Testing** |  |  |
| PyTest | MIT | Testing framework; pytest.org |
| GitHub Actions | Free* service | Continuous Integration; github.com/features/actions |
| Circle CI | Free* service | Continuous Integration; circleci.com |
| **Calibration** |  |  |
| PyBioNetFit [48*] | BSD | BNGL and SBML models |
| PyPESTO [49] | BSD | SBML and PETab support |
| PyDREAM [20*] | GPL | PySB interface |
| **Analysis & Visualization** |  |  |
| Matplotlib | PSF | Plotting library; matplotlib.org |
| Jupyter Notebooks | BSD | Computational notebooks; jupyter-notebook.readthedocs.io |
| PyVIPR [58*] | MIT | PySB, Tellurium interfaces |
| **Sharing and modification** |  |  |
| Github | Free* service | Code hosting and collaboration suite; github.com |
| Sphinx | BSD | Documentation framework; sphinx-doc.org |
| Readthedocs | Free*<br>service | Automated documentation compiler and hosting;<br>readthedocs.io |

## Acknowledgements

Funding was provided by the National Science Foundation (1411482 and 1942255 to C.F.L.) and the National Cancer Institute (U01CA215845 to C.F.L.).

